# Exploring protein-mediated compaction of DNA by coarse-grained simulations and unsupervised learning

**DOI:** 10.1101/2024.03.28.587201

**Authors:** Marjolein de Jager, Pauline J. Kolbeck, Willem Vanderlinden, Jan Lipfert, Laura Filion

## Abstract

Protein-DNA interactions and protein-mediated DNA compaction play key roles in a range of biological processes. The length scales typically involved in DNA bending, bridging, looping, and compaction (≥1 kbp) are challenging to address experimentally or by all-atom molecular dynamics simulations, making coarse-grained simulations a natural approach. Here we present a simple and generic coarse-grained model for the DNA-protein and protein-protein interactions, and investigate the role of the latter in the protein-induced compaction of DNA. Our approach models the DNA as a discrete worm-like chain. The proteins are treated in the grand-canonical ensemble and the protein-DNA binding strength is taken from experimental measurements. Protein-DNA interactions are modeled as an isotropic binding potential with an imposed binding valency, without specific assumptions about the binding geometry. To systematically and quantitatively classify DNA-protein complexes, we present an unsupervised machine learning pipeline that receives a large set of structural order parameters as input, reduces the dimensionality via principal component analysis, and groups the results using a Gaussian mixture model. We apply our method to recent data on the compaction of viral genome-length DNA by HIV integrase and we find that protein-protein interactions are critical to the formation of looped intermediate structures seen experimentally. Our methodology is broadly applicable to DNA-binding proteins and to protein-induced DNA compaction and provides a systematic and quantitative approach for analyzing their mesoscale complexes.

**SIGNIFICANCE:** DNA is central to the storage and transmission of genetic information and is frequently compacted and condensed by interactions with proteins. Their size and dynamic nature make the resulting complexes difficult to probe experimentally and by all-atom simulations. We present a simple coarse-grained model to explore ∼kbp DNA interacting with proteins of defined valency and concentration. Our analysis uses unsupervised learning to define conformational states of the DNA-protein complexes and pathways between them. We apply our simulations and analysis to the compaction of viral genome-length DNA by HIV integrase. We find that protein-protein interactions are critical to account for the experimentally observed intermediates and our simulated complexes are in good agreement with experimental observations.

## INTRODUCTION

DNA is central to the storage and transmission of genetic information, which critically involves a broad range of DNA-protein interactions. Both cellular and viral DNA are compacted by interactions with proteins, and recent evidence suggests that DNA often occupies cellular microenvironments or subcompartments, i.e. where DNA-protein interactions create condensates and membrane-less organelles (1–10). It has been shown that DNA-protein interactions are sufficient to compact DNA and create defined clusters (11–13). For example, vaccinia topoisomerase IB (vTopIB) was found to induce the formation of DNA-protein filaments at low protein-to-DNA ratios, by creating bridges between two segments of a single DNA molecule, and the formation of DNA-protein clusters of multiple DNA molecules at high protein-to-DNA ratios (14). Recent work has highlighted that DNA bridging can explain the compaction of DNA by SMC cohesin complexes in a phase diagram with an extended and a compacted phase (15). Similarly, DNA-protein interactions drive condensation involved in DNA repair (16) and in the compaction of mitochondrial DNA in nucleoids (17, 18).

As DNA looping and bridging –and ultimately compaction and clustering–typically involve length scales exceeding the DNA bending persistence length (40-50 nm, corresponding to ≈ 120-150 base pairs), characterizing the resulting mesoscale structures, either at high-resolution experimentally (19, 20) or by all-atom molecular dynamics simulations (21–24), becomes a challenging endeavor. Consequently, coarse-grained simulations can offer a highly complementary view, that can test mechanisms and provide microscopic insights not available directly from experiments, in the spirit of a computational microscope (25, 26). Coarse-grained simulations have provided many insights into DNA topology and dynamics (27–29), and coarse-grained simulations of simple DNA-protein models have contributed to our understanding of the formation of protein bridges and the resulting bridging-induced compaction (30–35). As a more specific example, coarsegrained models (36) explained the liquid droplet formation of heterochromatin due to heterochromatin protein 1 (37, 38).

In a recent experimental work, we observed DNA compaction by HIV integrase (IN) via a “rosette” intermediate (i.e. a central nucleo-protein core with extruding DNA loops), and introduced a coarse-grained model to explain this behavior (39). Here, we present this model in detail, as well as the analysis method that we used to understand the formation of DNA-protein complexes. Our coarse-grained model consists of DNA interacting with proteins, where both protein-DNA as well as protein-protein interactions are tuned to match experimental observations from Ref. (39). Specifically, the DNA is represented by a discrete worm-like chain, while proteins are treated as simple spherical particles. Our model is generic and can be readily extended to other proteins, as it reduces the protein-DNA interactions to a simple isotropic pair potential with a defined binding valency, without making specific assumptions about the binding geometry. The protein-DNA binding strength is taken from experimental measurements.

In order to characterize the effect of protein-protein interactions on the protein-mediated compaction of DNA, we present an unsupervised machine learning pipeline to systematically and quantitatively classify DNA-protein complexes: first, we define a set of structural parameters, next we reduce the dimensionality via principal component analysis, and finally divide the conformations into distinct groups with a Gaussian mixture model. We apply the approach to the IN-DNA compaction data (39), and find that protein-protein interactions are critical to the formation of looped intermediated structures (rosettes) seen experimentally. We expect our methodology to be widely applicable to DNA-binding proteins and to protein-induced DNA compaction.

## METHODS

We simulate the DNA-protein systems using Monte Carlo (MC) simulations of coarse-grained models. In this section, we introduce the specific representations of both the DNA and the proteins, and explain how we determine the binding strength between the proteins and DNA to match with experiments (39).

### Coarse-grained model for DNA

To model the double-stranded DNA, we use the common discrete worm-like chain model (WLC) (11, 15, 33, 40–43) and perform MC simulations of a double-stranded DNA in the canonical ensemble (44). This model is also frequently referred to as the beads-on-a-string model, as the DNA is treated as a string of *N* beads of diameter σ connected via finitely extensible nonlinear elastic (FENE) springs. The springs are described by the potential

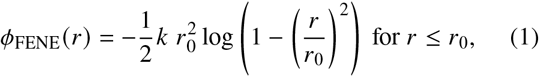

and infinity otherwise. Here, *k* = 21*k*_*B*_*T* /σ^2^ is the spring constant (15), *r* is the center-to-center distance between two beads, and *r*_0_ = 1.5σ is the maximum extension of the springs. Excluded volume interactions between the DNA beads are included via the Weeks-Chandler-Andersen (WCA) potential

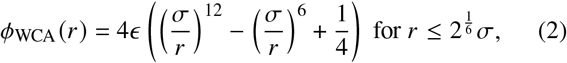

and zero otherwise. The interaction strength *ϵ* is set to 0.7*k* _*B*_*T* (15). The stiffness of the DNA chain is included via a bending potential

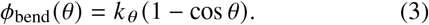

Here, *k* _*θ*_ is the spring constant for bending, and *θ* is the angle between successive springs. In order to imitate the experimental DNA, which has a persistence length of approximately 40 nm, we use DNA beads of σ = 4 nm and obtain the desired persistence length of 10σ by taking *k* _*θ*_ = 11*k*_*B*_*T*, see Supplemental Material.

In this work, we mainly consider four lengths of DNA: 150, 289, 408, and 774 beads, which correspond to roughly 1.8, 3.4, 4.8, and 9.1 kilo-base pairs (kbp). The latter three lengths correspond to DNA constructs used in experiments probing DNA interactions with HIV integrase (39), to enable qualitative comparison of our simulation results to experiments. Depending on the length of the worm-like chain, proper equilibration may require a significant amount of time. Hence, to speed up equilibration, each chain is initialized as a random walk with a fixed step size of σ and with bending angles evenly distributed between 0 *≤ θ ≤ θ*_max_. The maximum bending angle *θ*_max_ is set to 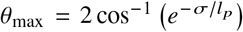 such that the initial chain has the approximate desired persistence length (42), i.e. *l* _*p*_ = 10σ. We equilibrate each DNA chain for at least 5 10^7^ MC cycles, which corresponds to roughly 5 10^5^ free diffusion times of a single, isolated DNA bead. Note that, on average, each DNA bead undergoes one trial move per MC cycle (44). During such a trial move the DNA bead is displaced by a random vector, whose individual *x*-*y*-, and *z*-components are drawn from a uniform distribution between ±0.12σ. We validate our model for DNA by reproducing the theoretical radius of gyration for a worm-like chain, see Supplemental Material.

### Coarse-grained model for proteins

Generally in DNA-protein systems, the proteins have a defined number of binding sites for DNA. For these kinds of systems, a patchy particle model is often used to simulate the coarse-grained multimer, see e.g. Refs. (11, 15, 35, 41, 45–47). However, as the exact geometry of the binding sites is often either unknown or poorly defined due to conformational flexibility of the protein, we decided against the use of a patchy particle model to prevent any potential restrictions of the patch geometry on the compaction of DNA. Instead, we choose to reduce the complex structure of the protein to a single protein bead with an isotropic binding potential and restrict the binding valence of the protein to *n*_max_ DNA beads (Fig. 1). Similar valence models have frequently been used to approximate patchy particles, see e.g. Refs. (48, 49). In this section we explain the mechanism behind this valence restriction.

**Figure 1:**
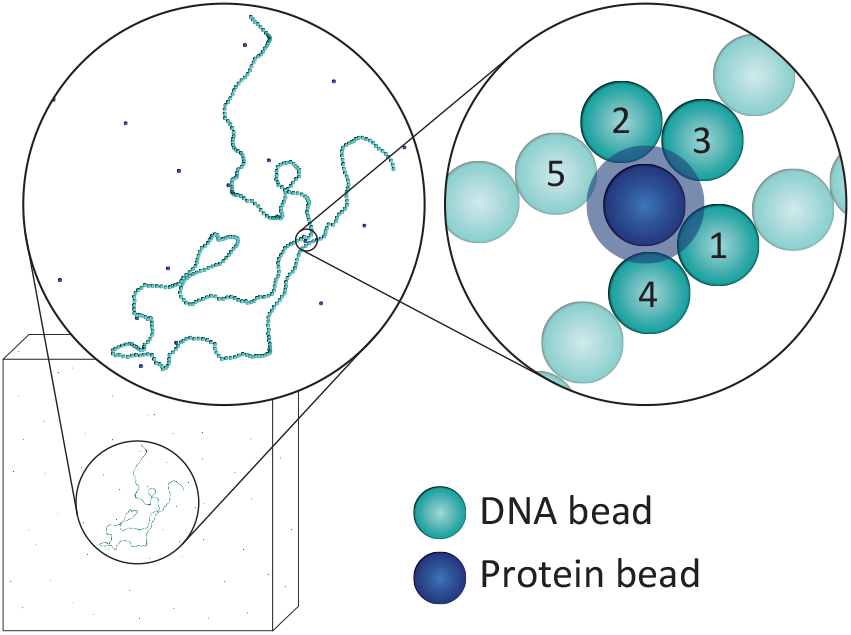
Schematic representation of the coarse-grained DNA-protein system. The zoom in shows the protein – with its isotropic binding potential – binding only the four nearest DNA beads.

For HIV integrase, which is our main focus here, the active complex binding to DNA is thought to be a tetramer with four binding sites. Furthermore, the binding footprint was experimentally determined to be 12 bp (39), which corresponds roughly to one tenth of the persistence length of DNA. Hence, we model the proteins as spheres of diameter σ, same as the DNA beads. However, one could easily simulate smaller or larger proteins. We ensure that the binding between the protein and DNA is short ranged, i.e. with an effective width of approximately 0.5 nm (50), by using the 18-36 Lennard-Jones (LJ) potential

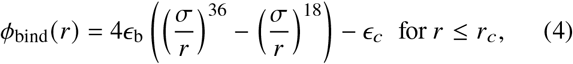

and zero otherwise. Here, *ϵ*_b_ is the binding strength, *r*_*c*_ = 1.4σ is the distance of truncation and *ϵ*_*c*_ is the energy shift such that the potential is zero at *r* = *r*_*c*_. The valence restriction is established by introducing a bond swapping potential

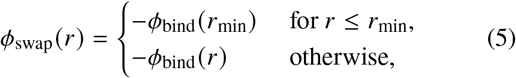

where *r*_min_ is the minimum of the binding potential. This potential is inspired by the one of Sciortino (51), and provides the possibility of freely swapping between (potential) bonds, i.e. without any additional energy cost or gain, while preserving both the condition of detailed balance and the excluded volume interactions that occur for *r < r*_min_. To illustrate how this works in general, assume that one wants to restrict the maximum number of bonds per protein to *n*_max_. For each protein, one then wants to take only the energy gain of the *n*_max_ shortest bonds into account. To accomplish this, we first find all DNA beads within the distance *r*_*c*_ of a protein *i* and sort them according to their center-to-center distance. We then compute the total binding energy of protein *i* using

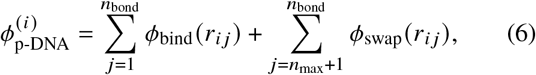

where 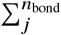 is the sum over the sorted list of *n*_bond_ DNA beads *j* with *r*_*i j*_ *< r*_*c*_. In this work, we use *n*_max_ = 4 to match with the four binding sites of the active integrase complex (39).

Lastly, we need to define the protein-protein interactions. In this work, we consider two cases for the protein-protein interaction. The first is a purely excluded volume interaction, which is included via the WCA potential with *ϵ* = 0.7*k* _*B*_*T*, same as for the DNA beads (Eq. 2). Secondly, we consider the possibility of some form of mutual attraction between the proteins in addition to the excluded volume interactions. This is realized by changing the protein-protein interactions from the WCA potential into the regular 6-12 LJ potential with attraction strength *ϵ*_pp_. The LJ potential is truncated and shifted at *r* = 3σ. We explore a couple of attraction strengths to find out what best fits the experimental observations.

### Protein-DNA binding strength

The key parameter describing the protein-DNA interactions is the binding strength *ϵ*_b_. Frequently, this binding energy has been treated as a free parameter in simulations of protein-DNA interactions, or it is estimated roughly from affinity measurements. Here, however, we use the binding strength taken from experimental measurements performed in Ref. (39). In order to obtain the protein-DNA binding strength, they used AFM images of short double-stranded DNA constructs in the presence of relative low concentrations of protein. By counting the number of protein molecules bound to the DNA constructs the binding probability can be directly determined, and is in turn compared quantitatively to simulations to obtain *ϵ*_b_. Note that the binding probability is computed by dividing the total number of bound protein molecules by the total number of DNA constructs considered and the number of possible binding sites on a DNA construct (assuming a binding footprint of 12 bp). The short DNA constructs and low IN concentration ensure that both multiple IN copies overlapping and protein-protein interactions are negligible. The binding probability increases monotonously with increasing binding strength with a crossover between simulations and experiments at a binding strength between 4.5*k* _*B*_*T* and 5.0*k* _*B*_*T* (39). We, therefore, take *ϵ*_b_ = 5*k* _*B*_*T* in this work.

Note that, for the experiments reported in Ref. (39), the integrase concentrations are expressed in terms of the monomer concentration, [IN]. However, since the complexes actually binding to the DNA are thought to be tetramers consisting of four integrase proteins, the protein concentrations in the simulations need to be adjusted to directly compare the experiments. Hence, we will express the protein concentration in terms of the monomer concentration throughout this work. Yet, keep in mind that the simulated protein beads, which represent the integrase tetramers, will have a concentration equivalent to one-fourth of the monomer concentration.

### Semi-grand canonical simulations to model DNA-protein mixtures

In our simulations, the DNA is treated in the canonical ensemble. However, since a substantial number of proteins can be involved in the protein-induced compaction of DNA, treating the proteins in the canonical ensemble can result in a significant depletion of the free proteins as our simulations box is not much larger than the DNA chain. Hence, to ensure that the concentration of free protein is not depleted in the simulations, we will simulate the protein in the grand-canonical ensemble. On top of the regular trial moves of a canonical ensemble, in the grand-canonical ensemble one needs trial insertions and removals of proteins (44). In order to prevent these insertions and removals from disturbing the DNA dynamics, and allow us to explore the intermediate structures during compaction, we restrict the part of the the simulation box where we perform insertion and deletion moves. Specifically, we take a spherical volume of radius *R* around the center of mass of the DNA chain, and prohibit the insertion or removal of integrases in this volume. The radius *R* is taken as the distance to the furthest DNA bead plus an additional padding of 5*r*_*c*_. Mirroring the experimental conditions, we will consider protein concentrations in the nanomolar to micromolar range. In the Supplemental Material, we provide more details on the semi-grand canonical simulations, and demonstrate that the concentration of free protein is not depleted during the compaction of DNA.

## RESULTS AND DISCUSSION

We examine the protein-induced compaction of DNA by performing a large set of simulations for the different protein-protein interactions (i.e. attractive and non-attractive) and for a large range of protein concentrations. For each protein-protein interaction and each protein concentration, we perform 10-12 independent simulations. To initialize the combined DNA-protein systems, we use an equilibrated DNA chain and insert the desired concentration of protein at random positions in the box. After equilibrating the total system for 10^6^ MC cycles with the DNA-protein interaction turned off, we gradually turn on the DNA-protein interaction within 100 MC cycles, and simulate for a maximum of another 5 · 10^8^ MC cycles.

During these simulations of the DNA-protein mixtures, we observe a range of different structures and conformations, depending on protein concentration and on the form and strength of their mutual interactions. Some examples of typical DNA-protein conformations are shown in Fig. 2*A-G*. Even though we can distinguish between some of these conformations by eye, it is difficult to classify all of them using a single order parameter like, e.g., the radius of gyration of the molecule. Hence, in order to systematically analyze and categorize the conformations formed in the simulations, we design an unsupervised machine learning pipeline.

**Figure 2:**
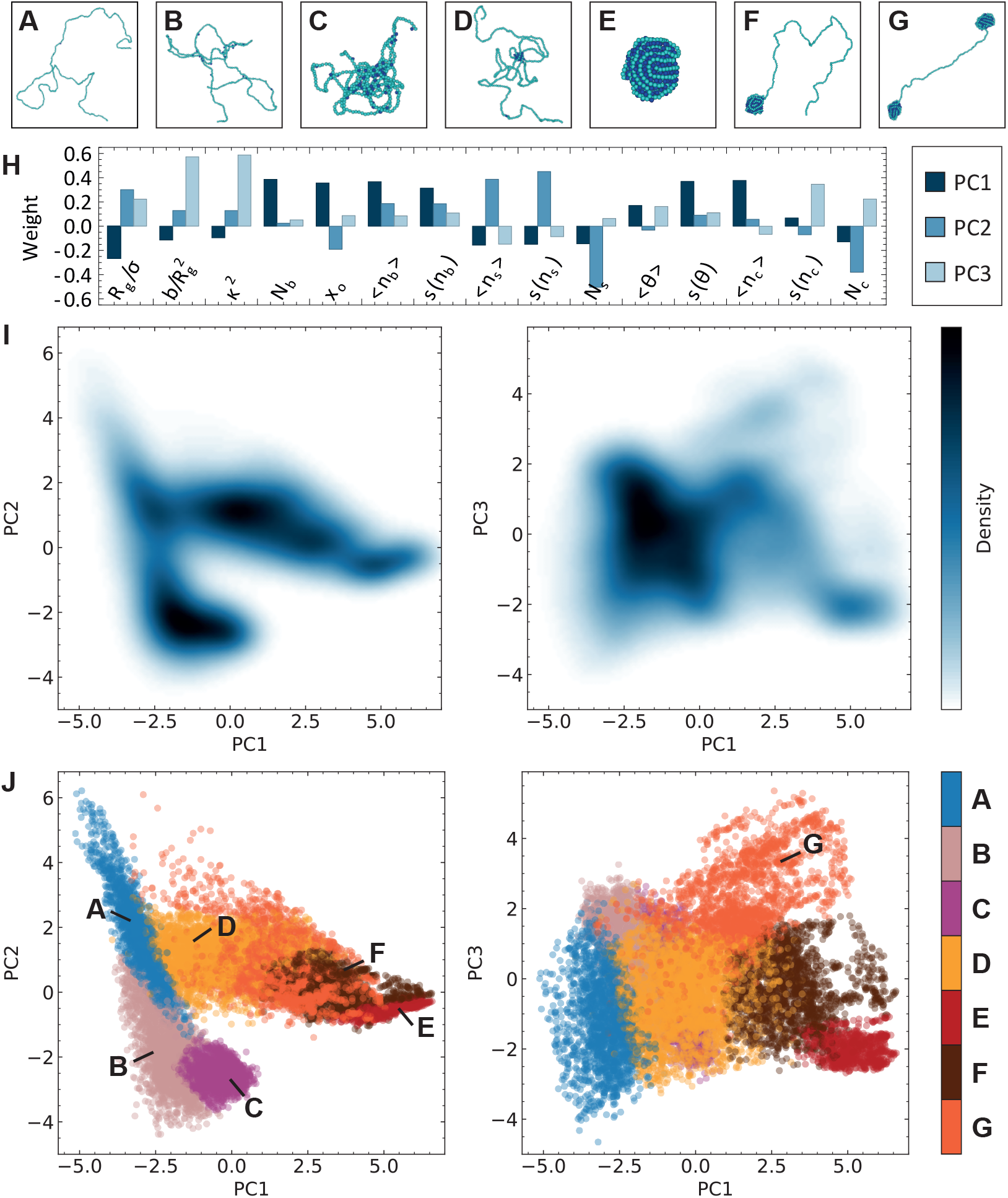
(*A-G*) Example conformations observed in simulations of DNA of 408 beads. Note that these are closeups and that the total 3D simulation box is many times larger. The DNA beads are indicated by light blue spheres and the proteins are indicated by dark blue spheres. For visualization purposes, the free proteins are not displayed. (*H*) The weight of each order parameter in the composition of the first three principal components (PCs). (*I*) Density distribution of the training dataset in the PC1-PC2 plane and the PC1-PC3 plane. (*J*) Grouping according to the Gaussian mixture model (GMM). The seven groups are indicated with different colors and they correspond to the typical conformations found in (*A-G*), i.e. (*A*) no compaction (blue), (*B*) bridging (pink), (*C*) bridging-induced compaction (purple), (*D*) rosette (yellow), (*E*) full compaction (red), (*F*) fully compacted complex with a bare tail (brown), and (*G*) multiple fully compacted complexes (orange).

The remainder of the Results and Discussion section is split into two. In the first part, we explain and set up the unsupervised machine learning pipeline for the classification of DNA-protein complexes. In the second part, we investigate the possible compaction pathways for the protein-mediated compaction of DNA by applying the trained machine learning pipeline to our simulations and classifying for each the structure of the DNA-protein complex as function of time.

### Unsupervised machine learned classification

In short, the unsupervised machine learning pipeline for the classification of protein-DNA complexes operates as follows. First it receives a multi-dimensional set of order parameters describing the geometrical characteristics of each conformation. It then applies a dimensionality reduction scheme which extracts its most important features. Lastly, a clustering algorithm identifies distinct groups of conformations in the resulting lower dimensional space. For the implementation of the dimensionality reduction and the grouping, we use the Python package scikit-learn (52). In this section, we explain the specifics of our classification approach in more detail.

#### Structural order parameters

To start, we first define a set of order parameters that capture the geometrical characteristics of each conformation. Note that for this initial selection of parameters, it is irrelevant whether parameters are independent or co-vary strongly. In the subsequent processing steps, these correlations are taken into account or can be removed if needed. So, while it is desirable to define parameters that capture a broad range of conformational features, it is not critical in our approach to a priori pick uncorrelated parameters. By looking at examples of different conformations (e.g. Fig. 2*A-G*), we compose a set of 15 parameters, which can be grouped into different categories:

- **Global conformation of the DNA chain**. The radius of gyration *R*_*g*_, the normalized asphericity 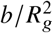, and the anisotropy *κ* of the DNA chain;
- **Bending angles of the DNA chain**. The average and standard deviation of the bending angle, *⟨θ*⟩ and s(*θ*);
- **Proteins bound**. The total number of proteins bound to DNA *N*_*b*_, the fraction of occupied DNA beads *x*_*o*_, and the average and standard deviation of the number of proteins bound per DNA bead, *⟨n*_*b*_⟩ and s(*n*_*b*_);
- **Bare DNA segments**. The total number of unoccupied DNA segments *N*_*s*_, and the average and standard deviation of the size of these unoccupied DNA segments, *⟨n*_*s*_⟩ and s(*n*_*s*_);
- **Protein clusters**. The total number of clusters of bound protein *N*_*c*_, and the average and standard deviation of the size of these clusters, *⟨n*_*c*_⟩ and s(*n*_*c*_).

Exact definitions of the different parameters can be found in the Supplemental Material.

#### Dimensionality reduction

To extract the important features of the 15-dimensional space of order parameters, we need to reduce its dimensionality. There are many options, both linear and nonlinear, for unsupervised dimensionality reduction, e.g. principal component analysis (PCA) (53–56), autoencoders (57–60), and manifold learning methods (61–64). For our problem, we found that PCA, despite being purely linear, robustly provides a satisfactory separation of the various states of compaction. To check for the effects of linearity, we have also confirmed that using a simple nonlinear neural-network-based autoencoder, like the one in Ref. (59), does not improve our ability to classify structures for this problem. Since PCA is computationally efficient, deterministic and parameter-free, we here choose to use it for the remainder of this manuscript.

It is important to note that, in general, dimensionality reduction schemes require a balanced dataset, consisting of a fairly equal representation of the various possible states, for the scheme to perform well. Hence, as a first test to demonstrate our classification method, we focus on DNA of 408 beads for which we compose a training set of nearly 15,000 configurations. This dataset contains roughly equal numbers of non-compacted configurations, configurations in different states of compaction mediated by mutual non-attractive proteins, and the same for compaction mediated by mutual attractive proteins with an attraction strength of 2.0*k* _*B*_*T*. We later explain how to extend this to include configurations of DNA of different lengths.

After constructing this balanced set of configurations, we normalize the distribution of each order parameter using standard scaling before feeding it to PCA. This ensures that each order parameter has an average value of zero and a variance of one, such that the variations in each parameter are treated as equally important. We determine the number of relevant principal components (PCs) by looking at the proportion of variance explained (PVE) of each PC. In this case, the first three PCs combined capture more than 75% of the total variance of the dataset. Further analysis on the PVE using the elbow method (65) also confirms the use of the first three PCs, see Supplemental Material.

The weight of each order parameter in the composition of the first three PCs is shown in Fig. 2*H*, and Fig. 2*I* shows the density distribution of the training dataset. We can already by eye distinguish some groups in this density distribution. For example, we see a distinct group on the top left of the PC1-PC2 plane from which two separate branches grow. The weights reveal that configurations in this group have a large radius of gyration and not many bound proteins; hence, this group most likely contains configurations of non-compacted DNA. Note that the weights also reveal that some order parameters are indeed highly correlated and therefore most likely redundant. For example, the asphericity and the anisotropy have very similar weights, as well as the average and standard deviation of the number of proteins bound per DNA bead. At this stage, one could spend some time sieving out the redundant order parameters. However, since the resulting classification was already sufficient in our case, we did not do this here.

#### Identifying the distinct DNA-protein conformations

To finish up the classification, we use a clustering algorithm to divide the three-dimensional distribution of PCs into distinct regions. As for the dimensionality reduction, there are many options for clustering (66), e.g. K-means, spectral clustering, and Gaussian mixture models. We find that, in our case, the Gaussian mixture model (GMM) provides a satisfactory distinction of the different groups in the PC landscape. We want to stress that the term “clustering” here means the classification of DNA-protein configurations into distinct groups with similar geometric features and should not be confused with the condensation or compaction of DNA-protein complexes which can also be called clustering.

To determine the number of groups, we first look for a minimum in the Bayesian information criterion (BIC) (67), which indicates how well a GMM fits the distribution while simultaneously penalizing on the number of groups to prevent overfitting. However, as an alternative criterion to safeguard against overfitting, we additionally look for an elbow in the clustering entropy (68)

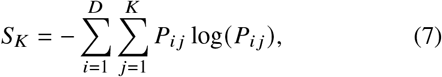

where *D* is the size of the dataset that needs to be grouped, *K* is the number of groups, and *P*_*i j*_ is the probability of data point *i* to belong to group *j*. We find that the optimum number of groups is seven, see Supplemental Material.

By definition, the GMM provides a “soft grouping”, i.e. the probability for a given configuration to belong to any of the groups. To turn this into a discrete grouping, we assign each configuration to the group it is most likely to belong to. Figure 2*J* shows the resulting grouping using seven groups for the GMM. Comparing Figs. 2*J* and 2*I*, we see that these groups nicely correspond to the groups visible in the density profile. Furthermore, by looking at various configurations belonging to these seven groups, we can identify them. Each group can be identified by one of the (deliberately chosen) example conformations depicted in Fig. 2*A-G*, i.e. (*A*) no compaction, (*B*) bridging, (*C*) bridging-induced compaction, (*D*) rosette, (*E*) full compaction, (*F*) fully compacted complex with a bare tail, and (*G*) multiple fully compacted complexes connected by bare segment(s) of DNA.

### Two pathways for the protein-mediated compaction of DNA

In order to investigate the effect of the protein-protein interactions on the time evolution of the protein-mediated compaction of DNA, we apply our trained classification method to our simulations of DNA of 408 beads. For each simulation, we classify at regular time intervals the structure of the DNA-protein complex such that we can easily interpret its time evolution. To demonstrate the strength of the machine learned classification, in Fig. 3*A,B* we show two typical simulation trajectories, the first for mutually non-attractive proteins (Fig. 3*A*) and the second for mutually attractive proteins (Fig. 3*B*). We clearly see that the two simulations follow two very different paths in the landscape of the principal components. Moreover, taking the classification of the GMM into account, we find that these are even two completely separate pathways, i.e. a pathway that goes via bridging to bridging-induced compaction for non-attractive protein-protein interactions (blue-pink-purple classification) and a pathway that goes via rosette to full compaction for attractive protein-protein interactions (blue-yellow-brown-red classification).1

**Figure 3:**
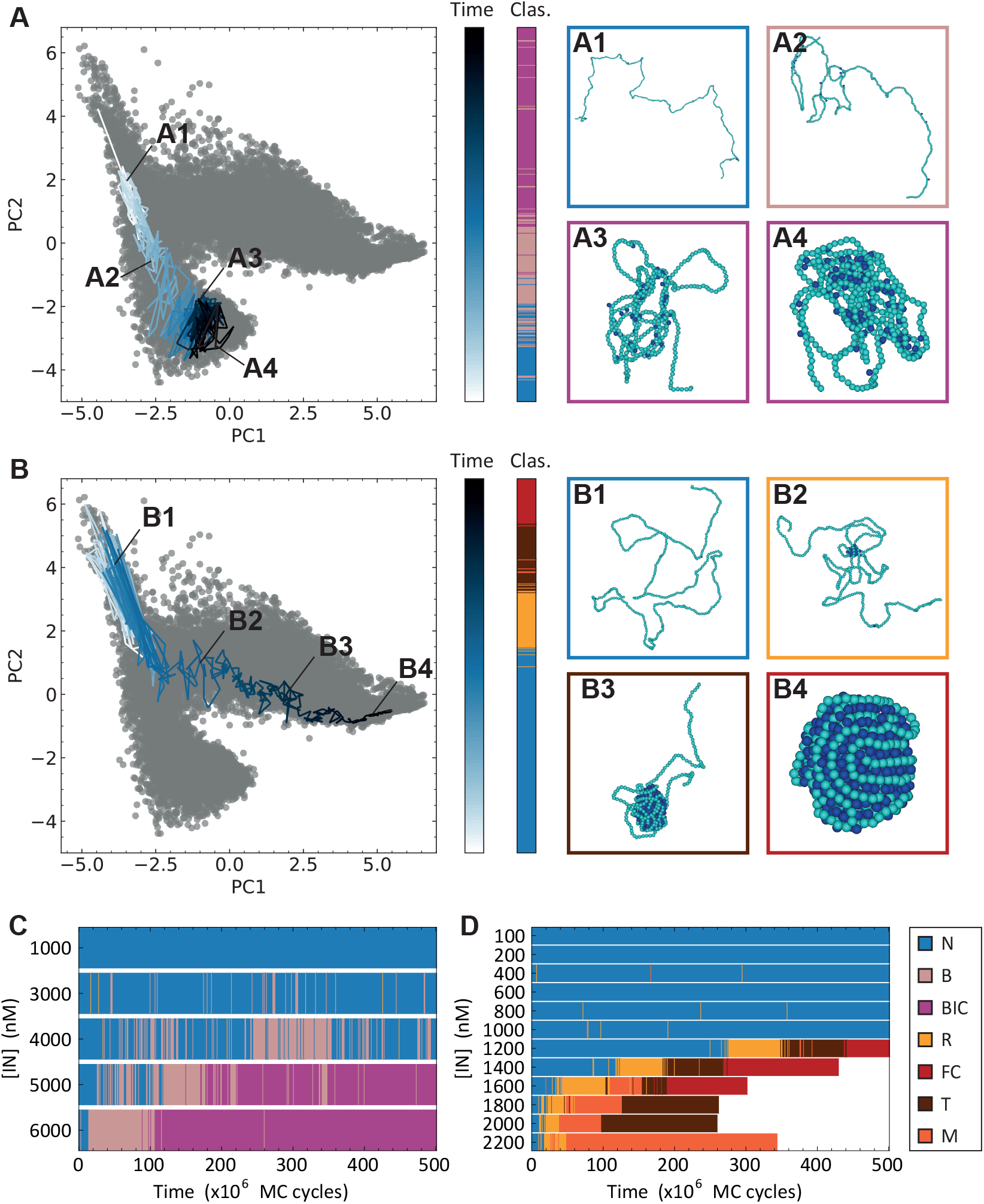
Typical simulations of DNA of 408 beads in (*A,C*) a system with non-attractive protein-protein interactions and (*B,D*) a system with attractive protein-protein interactions of strength *βϵ*_pp_ = 2.0. (*A,B*) Show, for a protein concentration of, respectively, [IN] = 5000 nM and [IN] = 1200 nM, the simulation trajectories on top of PC1-PC2 distribution of the training dataset. The trajectories are colored with a blue gradient indicating the simulation time (first colorbar) and the second colorbar is the classification according to the GMM. The figure blocks (*A1-A4*) and (*B1-B4*) each show four characteristic configurations during the simulation. (*C,D*) Time evolution of the classification as a function of protein concentration. Each bar represents the most typical run for each protein concentration studied. The seven options for the classification are (see Fig. 2): no compaction (N), bridging (B), bridging-induced compaction (BIC), rosette (R), full compaction (FC), fully compacted complex with a bare tail (T), and multiple fully compacted complexes (M). When a bar ends in white it means the simulation was terminated early as the most interesting behavior had already happened.

To establish that this is not accidental of these two specific runs or protein concentrations, we examine the time evolution of the classification of all 10 to 12 simulations per protein concentration and protein-protein interaction. These simulations indeed confirm that there is no pathway between the bridging and rosette states. The compaction pathway is fully determined by the protein-protein interaction. To illustrate that this result is independent of the protein concentration, we select the most typical simulation per concentration and show the associated time evolution of the classification in Fig. 3*C,D*. These trajectories are selected based on the inverse of the time the system needs to start (irreversible) compaction and the time spent in either the bridging or rosette state. A trajectory is coined “most typical” when these (inverse) times best match with the average (inverse) times for the system at that protein concentration. One can clearly see that there is no crossover between the pathway of bridging-induced compaction (oberved in 3*C*) and the pathway of rosette to full compaction (observed in 3*D*).

Furthermore, it is noteworthy that from these simulations we can conclude that protein-protein attractions promote DNA compaction; Fig. 3*C,D* clearly shows that DNA compaction in systems with 2*k* _*B*_*T* protein-protein attraction sets in at lower protein concentrations than in systems without protein-protein attraction. Although this is not surprising – protein-protein attractions naturally lead to a higher affinity to compact –, it is interesting to explore the effect of the attraction strength on the onset of DNA compaction.

#### Role of DNA length and protein-protein attraction strength

To explore the effect of protein-protein attraction strength in the protein-induced compaction of DNA, we focus on attraction strengths of 1.5*k* _*B*_*T* and 2.0*k* _*B*_*T* and protein concentrations from 100 nM to 2400 nM. Additionally, we investigate the role of DNA length by considering DNA of 150, 289, 408, and 774 beads. Importantly, given the dependence of some of the structural order parameters used as input in the classification on DNA length (e.g. the number of bound proteins or the length and number of bare DNA segments), we need to adjust our classification to accommodate different DNA lengths. In order to use one pipeline for the classification of systems with different DNA lengths, we first obtain a balanced training dataset for each DNA length which we then normalize separately from each other. This ensures that, even though the important features differ in absolute values, they are treated roughly as equal in each normalized dataset. Note that we were able to do this as no new or distinctly different structures emerged as a result of varying DNA length and protein-protein attraction strength.

Next, we combine these separately normalized sets and obtain a new classification by retraining both the PCA and GMM on the combined dataset. To demonstrate that the new classification is indeed able to identify conformations like the rosette conformation for the different DNA lengths, we show a short time series of a typical trajectory for DNA of 150, 289, and 774 beads in Fig. 4. Even though the rosette conformation, for example, looks undeniably different for different DNA lengths, the classification is able to identify it as the same conformation for all DNA lengths.

**Figure 4:**
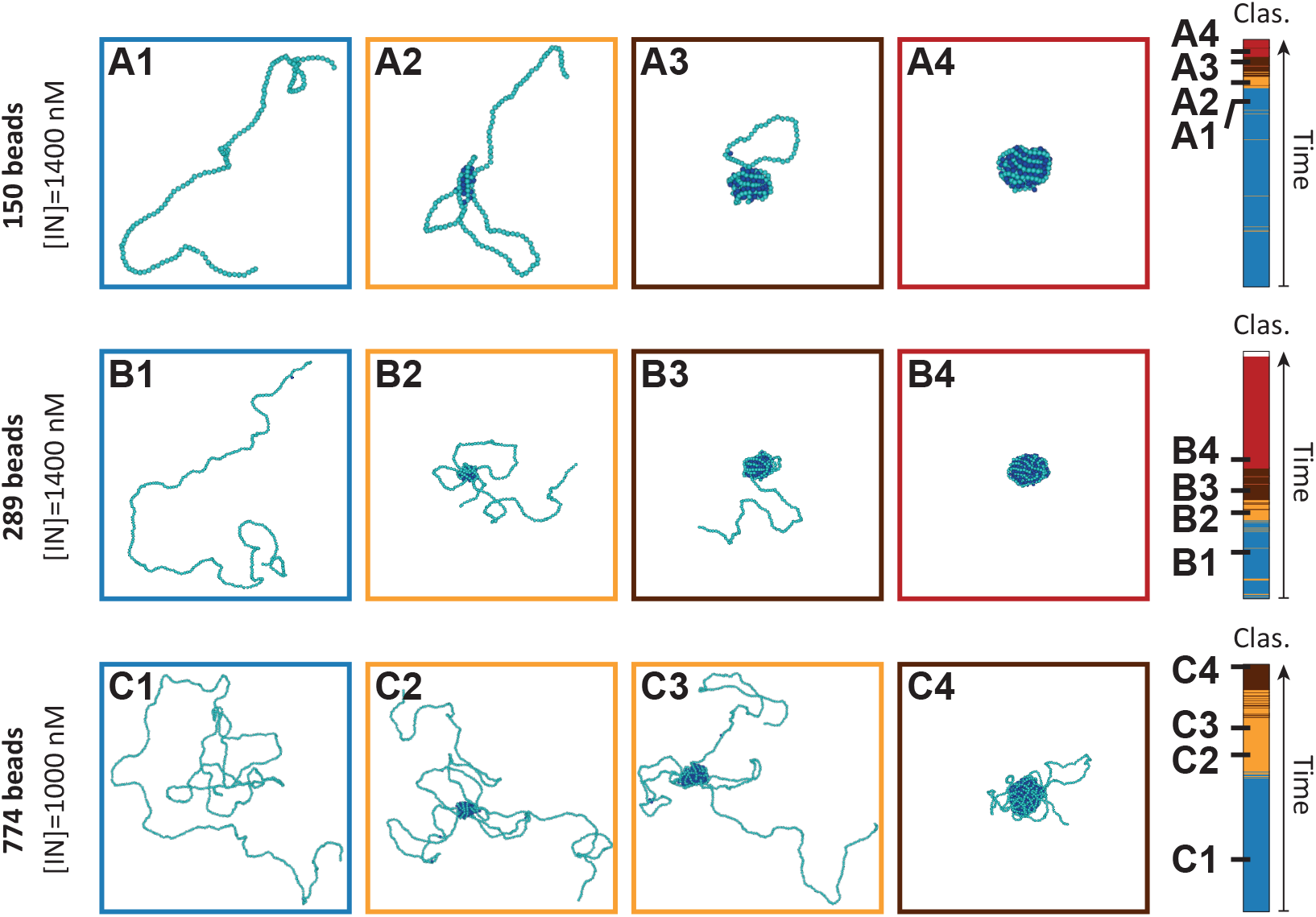
Short time series of a typical simulation of DNA of (*A1-A4*) 150 beads, (*B1-B4*) 289 beads, and (*C1-C4*) 408 beads. In all cases the DNA is in solution with mutual attractive proteins (*βϵ*_pp_ = 2.0). The colorbars indicate the classification as a function of time, see Fig. 3.

Using the newly trained pipeline, we classify all 12 simulations per DNA length and protein-protein attraction strength. To illustrate the influence of both DNA length and protein-protein attraction strength on the protein-mediated compaction of DNA, we again select the most typical simulation per concentration and system, and show the associated time evolution of the classification in Fig. 5. Note that, as mentioned before, these typical trajectories are selected based on the inverse of the time the system needs to start (irreversible) compaction and the time spent in the rosette state. There are four key observations that can be deduced from Fig. 5. First, we find that all trajectories leading to compaction go through a transient rosette state, independent of the DNA length or protein concentration. Second, it is evident that stronger protein-protein attractions facilitate DNA compaction at lower protein concentrations for all investigated DNA lengths. Similarly, we see that an increase of DNA length also enables compaction at lower protein concentrations. Lastly, we observe that the time spent in a rosette conformation (indicated in yellow) increases with the DNA length.

**Figure 5:**
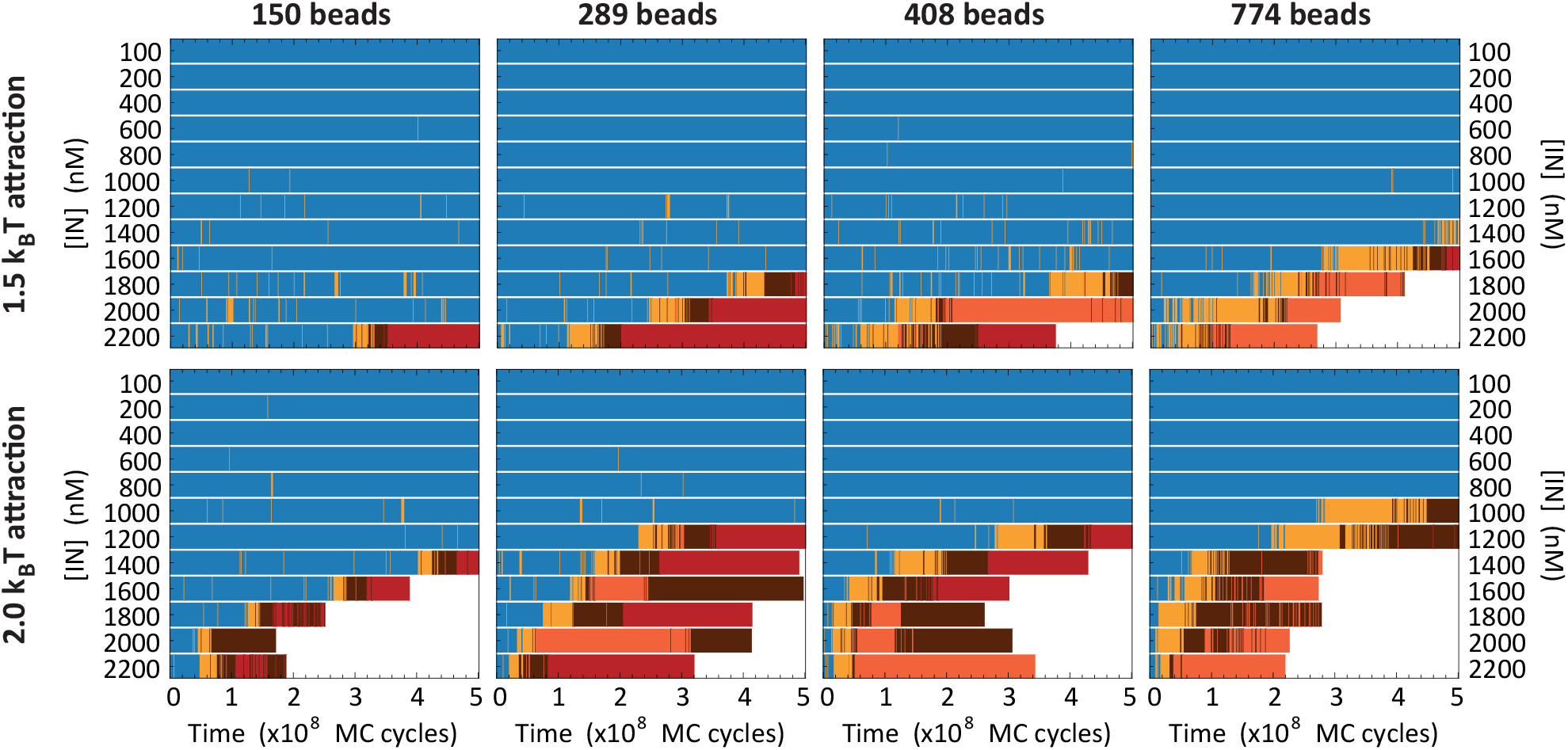
The most typical runs for a range of protein concentrations for the four DNA lengths studied. The top row gives the results for a mutual protein interaction with attraction strength *βϵ*_pp_ = 1.5 and the bottom row *βϵ*_pp_ = 2.0. See Fig. 3 for the interpretation of the classification that the colors of the bars represent.

In order to translate some of these observations into quantitative measures, we compute the rate of compaction as a function of protein concentration for the different systems. Assuming that the start of compaction in our simulations is a rare event which has a censored exponential distribution, i.e. not necessarily all simulations managed to compact, we can approximate the rate *λ* with (69)

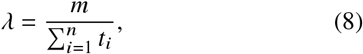

where the sum is taken over the *n* = 12 simulations performed for a specific system, *m* is the number of simulations in which the DNA managed to compact, and *t*_*i*_ is the time of simulation *i* at which (irreversible) compaction starts. Practically, *t*_*i*_ is the last instance of a conformation classified as not compacted (indicated by blue in Fig. 5). Note that the sum over all *n* = 12 simulations also includes the time of the simulations in which the DNA did not manage to compact. The resulting rates are given in Fig. 6. Here we have normalized the rates with the DNA length, such that a rough collapse onto a master curve for both protein-protein attraction strengths is revealed. The trends of these master curves suggest that the compaction of DNA is an activating event with a rate given by

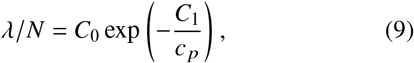

where *c* _*p*_ is the protein concentration and *C*_0_ and *C*_1_ are parameters which depend on the protein-protein attraction strength.

**Figure 6:**
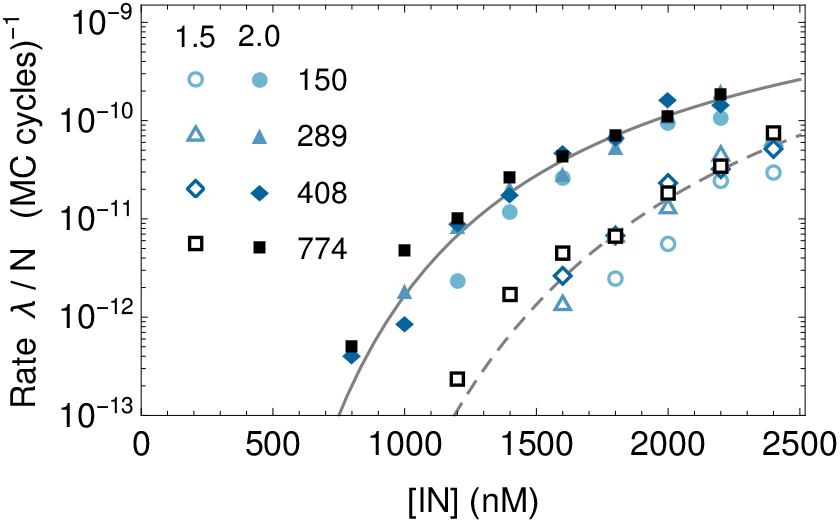
The rate of compaction divided by the DNA length as a function of the protein concentration. The shapes of the markers indicate the four different lengths of DNA, and the two different attraction strengths *βϵ*_pp_ = 1.5 and *βϵ*_pp_ = 2.0 are indicated by the open and closed markers, respectively. The lines indicate fits to Eq. 9.

Note that, although the translation to real time is not as direct as for e.g. molecular dynamics simulations, the obtained rates can still be roughly related to real time given that the long-time diffusion of an isolated DNA bead is ∼ 100 MC cycles in our simulations. Moreover, the relative trends obtained from comparing different systems give legitimate insights into the workings of protein-mediated compaction of DNA.

## CONCLUSION

To conclude, we presented a simple coarse-grained model for DNA-protein mixtures that treats the protein-DNA interactions as an isotropic binding potential with an imposed binding valency without specific assumptions about the binding geometry. This approach for the protein-DNA binding presents a generic solution for the many cases in which the exact geometry of the binding sites is either unknown or poorly defined due to conformational flexibility of the protein. Additionally, we designed a simple, fast, and effective unsupervised machine learning model for the classification of the DNA-protein complexes into different conformational states. We applied our model to recent data on the compaction of viral genome-length DNA by HIV integrase and we find that protein-protein attractions are critical to the formation of looped intermediated structures (“rosettes”) observed experimentally. Not only is our model applicable to a broad range of different protein and nucleic acid systems, the additional unsupervised learning method for the classification of the intricate complexes formed in such systems can offer key insights into the variety of conformational states and their formation pathways.

## ACKNOWLEDGMENTS

We would like to thank Frank Smallenburg and Rinske Alkemade for many useful discussions. We acknowledge funding from the Vidi research program with project number VI.VIDI.192.102 which is financed by the Dutch Research Council (NWO) and by Utrecht University.

## Footnotes

1 Note that here we observe two clear pathways that we connect to attractive and non-attractive protein-protein interactions. However, note that this is only true for sufficiently large attractive strengths.

