## Supplemental Material for "Exploring protein-mediated compaction of DNA by coarse-grained simulations and unsupervised learning"

### I. JUSTIFICATION OF THE MODEL

#### A. Radius of gyration

In order to validate our coarse-grained model for double-stranded DNA, we compute the average radius of gyration as a function of the DNA length and compare it to the theoretical trend of the worm-like chain (WLC) model. We consider DNA of 150, 289, 408, and 774 beads of diameter $\sigma$. For each DNA length, we run 50 simulations of $10^8$ Monte Carlo (MC) cycles of individually equilibrated chains and measure the radius of gyration each $10^6$ MC cycles. The resulting average radius of gyration as a function of the DNA length is given in Fig. S1. The dashed line in this figure indicates the radius of gyration according to the WLC model, which is given by[1,2]

$$R_g^2 = \frac{Ll_p}{3} - l_p^2 + \frac{2l_p^3}{L} - \frac{2l_p^4}{L^2}\left(1 - e^{-L/l_p}\right), \tag{S1}$$

where $l_p$ is the persistence length, $L = Na$ is the contour length, with $N$ the number of monomers and $a$ the effective monomer length. For our system, $l_p = 10\sigma$ and $a = 0.974\sigma$. We see that the radius of gyration agrees well with the theoretical model. Additionally, they align with values reported from experiments on linear DNA[3].

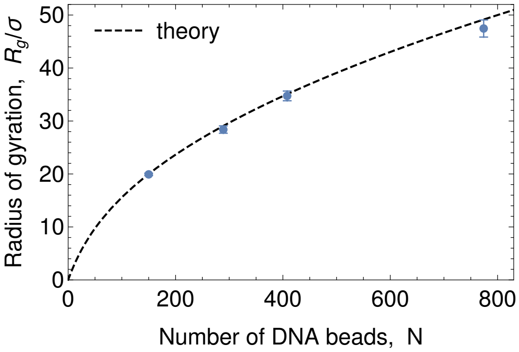


FIG. S1. The radius of gyration as a function of DNA length as measured in the simulations. The dashed line indicates the theoretical trend, Eq. (S1) with $l_p = 10\sigma$ and $a = 0.974\sigma$.

#### B. Grand-canonical ensemble of proteins

In the experiments, the relative concentration of integrase with respect to DNA is sufficiently high such that, even for the lowest integrase concentrations considered, the free integrase is not depleted as a result of the compaction of the DNA[4]. To replicate this condition in simulations, it is necessary to have a substantial reservoir of protein. One straightforward approach to achieving this is by using an exceptionally large simulation box. This, however, would significantly slow down the simulations. A more effective alternative is to establish an external protein reservoir in the form of a grand-canonical ensemble. We, therefore, chose to simulate the DNA in the canonical ensemble and the proteins in the grand-canonical ensemble. Note that, in order to directly compare with experiments, we express the protein concentration in terms of the monomer concentration, [IN]. This means that the protein beads representing the integrase tetramers are simulated at a concentration equivalent to one-fourth of the monomer concentration.

As explained in the main text, we obtain the grand-canonical ensemble via trial insertions and removals of proteins. To prevent any interference with the DNA-protein complex, we only apply these trial insertions and removals to the part of the system that does not contain the DNA chain. More specifically, this subsystem consists of the entire simulation box excluding a sphere of radius $R$ around the center of mass of the DNA chain. Following the book by

Frenkel and Smit[5], the trial insertions into and removals from this subsystem are accepted with the probabilities

$$P_{\text{insert}}(N_p \to N_p+1) = \min\left[1, \frac{zV}{N_p+1}\, e^{-\beta[U(N_p+1)-U(N_p)]}\right], \tag{S2}$$

$$P_{\text{remove}}(N_p \to N_p-1) = \min\left[1, \frac{N_p}{zV}\, e^{-\beta[U(N_p-1)-U(N_p)]}\right], \tag{S3}$$

where $N_p$ is the number of proteins in the subsystem before the trial insertion or removal, $V$ is the volume of the subsystem, $U(N_p)$ is the internal energy of the subsystem with $N_p$ particles, and $z = e^{\beta\mu}/\Lambda^3$ is the fugacity with $\mu$ the chemical potential. Since the protein concentrations considered in this work are extremely low, e.g. [IN] = 100nM corresponds to a number density of $\rho\sigma^3 \approx 10^{-6}$, the chemical potential can be approximated by that of an ideal gas, $\beta\mu_{\text{id}} = \ln(\rho\Lambda^3)$. Consequently, the fugacity is simply equal to the desired number density of the system. Hence, the fractions in Eqs. S2 and S3 can be viewed as ratios between the actual and desired number of proteins in the subsystem.

To demonstrate that the concentration of free protein is indeed not depleted during the compaction of DNA, Fig. S2 shows for a typical compaction event the number of total, free, and bound proteins in the system. One can clearly see that the number of free proteins fluctuates in a stable fashion around the target value, even when the DNA starts compacting at around $1.8 \cdot 10^8$ MC cycles. Note that for all simulations with an attractive protein-protein interaction we use a simulation box of $(350\sigma)^3$, and for all simulations with a non-attractive protein-protein interaction we use a simulation box of $(250\sigma)^3$. All simulations have periodic boundary conditions.

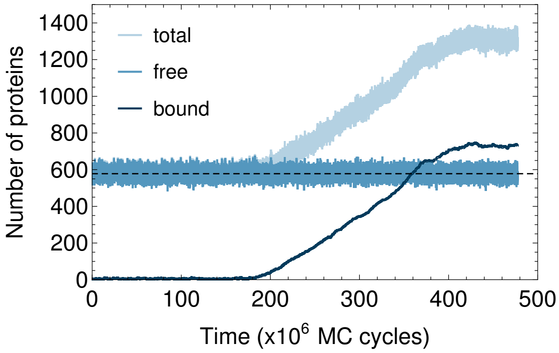


FIG. S2. The number of total, free, and bound proteins during a typical protein-induced compaction event of DNA. This particular event is of DNA of 408 beads in a system with attractive protein-protein interactions of strength $\beta\epsilon_{\text{pp}} = 2.0$ at concentration [IN]=1400 nM. The horizontal dashed line indicates the target number of free proteins.

### II. ADDITIONAL INFORMATION ON THE MACHINE LEARNED CLASSIFICATION

#### A. Input parameters

In the main text, we list the 15 structural order parameters that we use as input for our unsupervised machine learned classification. The definitions of the majority of these parameters speak for themselves. Here, we provide the exact definition for the parameters for which that is not directly clear.

The radius of gyration, asphericity, and anisotropy are all obtained from the eigenvalues of the gyration tensor of the DNA chain:

$$R_g^2 = \lambda_0 + \lambda_1 + \lambda_2, \quad b = \lambda_2 - \frac{1}{2}(\lambda_0 + \lambda_1), \quad \text{and} \quad c = \lambda_1 - \lambda_0, \tag{S4}$$

where the eigenvalues are ordered as $\lambda_0 \leq \lambda_1 \leq \lambda_2$. Here the gyration tensor is defined as

$$S_{mn} = \frac{1}{N}\sum_{i=1}^{N} r_m^{(i)} r_n^{(i)}, \tag{S5}$$

where $r_m^{(i)}$ is the $m$th coordinate of the position of the $i$th DNA bead. For simplicity, we only take DNA beads into account in the calculation of the gyration tensor and not any bound protein beads. However, including the bound protein in the calculation of the gyration did not significantly effect the final classification of the protein-DNA conformations.

Neighboring bound proteins are defined as a protein cluster when the cluster contains at least two bound proteins. Within this definition, two bound proteins belong to the same protein cluster when their center-to-center separation is less than $3.0\sigma$.

#### B. Principal component analysis

In the main text, we explain the use of principal component analysis (PCA) for the dimensionality reduction of the 15-dimensional space of order parameters. We explain that we decided to use the first three principal components (PCs) and disregard the others. Here, we provide further validation for this choice. The proportion of variance explained (PVE) by each PC reveals that the first three PCs account for 41%, 19%, and 16% of the variance, totalling to 76%, see Fig. S3. The subsequent PCs capture significantly less information, e.g. PC4 captures only 6.4%. This alone provides substantial support for our decision to use the first three PCs. However, we further confirm our choice by determining the location of the elbow in the PVE. For this we use L-method of Salvador and Chan[6]. The two most obvious candidates for the location of the elbow are at the third or the fourth PC. Comparing the linear fits for an elbow at PC3 (Fig. S3*A*) and for an elbow at PC4 (Fig. S3*B*), we discover that the elbow at PC3 has the lowest root means squared error (RMSE). Hence, we conclude that three PCs is the best choice for the dimensionality reduction.

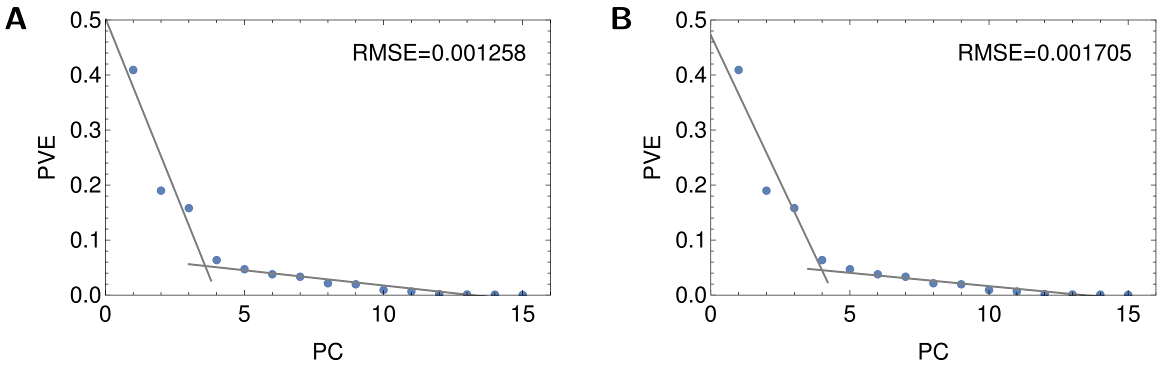


FIG. S3. The proportion of variance explained (PVE) by each principal component (PC). The gray lines represent linear fits indicating a potential elbow at (*A*) 3 PCs and (*B*) 4 PCs. The total root mean squared error (RMSE) of the fits reveals that the elbow is located at the third PC.

#### C. Gaussian mixture model

In the main text (Fig. 2), we present the density distribution of the training dataset within the principal component (PC) planes, and highlight the presence of distinct groups within this distribution. We further describe the use of a Gaussian mixture model (GMM) for a unsupervised grouping of this space and, in particular, explain how we come to the choice of seven groups. Here, we provide the additional information behind this choice. In Fig. S4*A*, we show the Bayesian information criterion (BIC)[7] as a function of the number of groups. One generally searches for a minimum in the BIC, as this number tells you how well a GMM fits the distribution while simultaneously penalizing on the number of groups to prevent overfitting. However, since this is not a foolproof method against overfitting, especially when a distribution contains (partially) overlapping groups, we additionally look at for an elbow in the clustering entropy[8]. Figure S4*B* reveals the existence of two elbows in the clustering entropy: one at four groups and another at seven groups. Based on the density distribution (Fig. 2*I* main paper), we can dismiss the option of four groups, as this distribution clearly shows more than four distinct groups. We, thus, end up with seven groups for the GMM. To, nonetheless, gain insight into what a separation into four groups might look like, Fig. S5 shows the resulting separation into four groups. We see that the GMM still manages to separate the bridging conformations from the other forms of compaction; however, with four groups one loses the distinction between the partially compacted conformations, i.e. rosette, fully compacted with bare tail, and multiple fully compacted complexes.

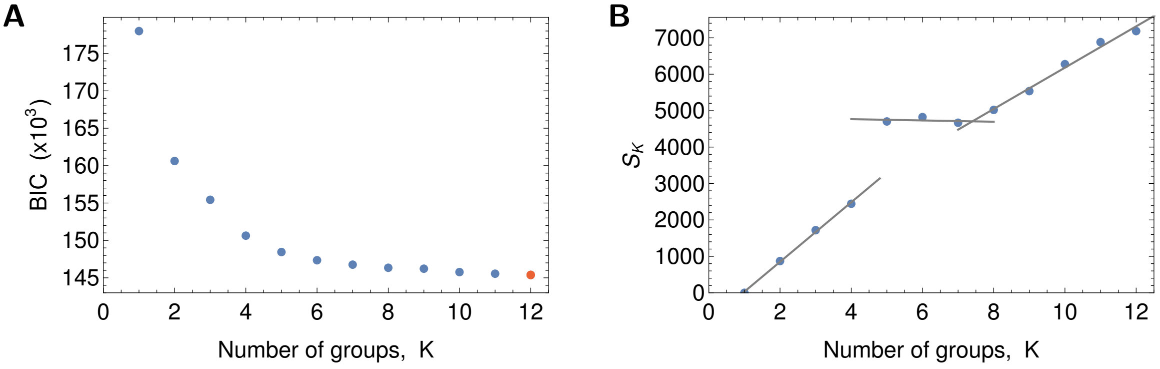


FIG. S4. (*A*) The Bayesian information criterion (BIC) as a function of the number of groups for the GMM. The minimum is highlighted in red. (*B*) The clustering entropy $S_K$ as a function of the number of groups. The gray lines represent linear fits indicating elbows at 4 and 7 groups.

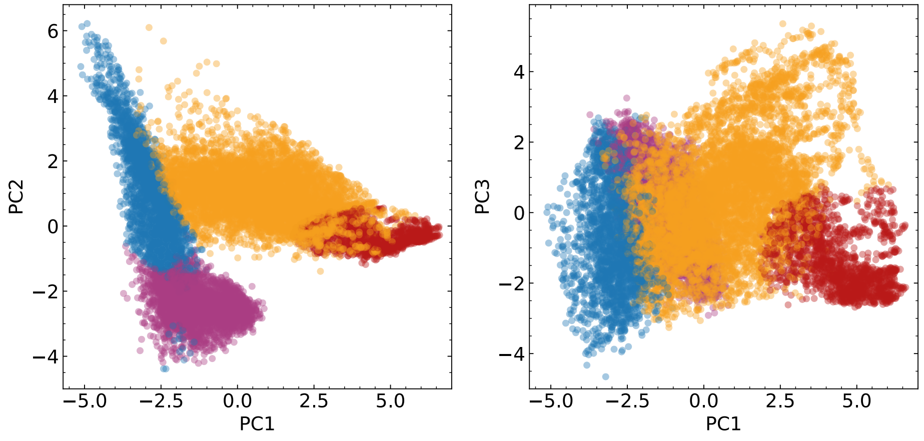


FIG. S5. Grouping of the distribution of the first three principal components (PCs) according to a Gaussian mixture model (GMM) with four groups.

### D. Additional example trajectory

In the main text (Fig. 3), we provide two examples of typical trajectories in the landscape of the principal components for the protein-induced compaction of DNA of 408 beads. Here, we show an additional trajectory to illustrate the need of the third principal component (PC3). Figure S6 shows a typical trajectory that leads to the protein-induced compaction into two separate, fully compacted DNA-protein complexes. The figure clearly illustrates need of PC3 to identify the compaction into separate DNA-protein complexes (classified as orange).

[1] J. B. Hays, M. E. Magar, and B. H. Zimm, Biopolymers **8**, 531 (1969).
[2] H. Benoit and P. Doty, J. Phys. Chem. **57**, 958 (1953).
[3] R. M. Robertson, S. Laib, and D. E. Smith, Proc. Natl. Acad. Sci. **103**, 7310 (2006).
[4] P. J. Kolbeck, M. de Jager, M. Gallano, T. Brouns, B. Bekaert, W. Frederickx, S. F. Konrad, S. V. Belle, F. Christ, S. D. Feyter, Z. Debyser, L. Filion, J. Lipfert, and W. Vanderlinden, bioRxiv (2024), 10.1101/2024.03.15.585256, https://www.biorxiv.org/content/early/2024/03/17/2024.03.15.585256.full.pdf.
[5] D. Frenkel and B. Smit, *Understanding Molecular Simulation: From Algorithms to Applications*, 2nd ed. (Academic Press, San Diego, 2002).
[6] S. Salvador and P. Chan, in *16th IEEE international conference on tools with artificial intelligence* (IEEE, 2004) pp. 576–584.
[7] G. Schwarz, Ann. Stat. , 461 (1978).
[8] J.-P. Baudry, A. E. Raftery, G. Celeux, K. Lo, and R. Gottardo, Journal of computational and graphical statistics **19**, 332 (2010).

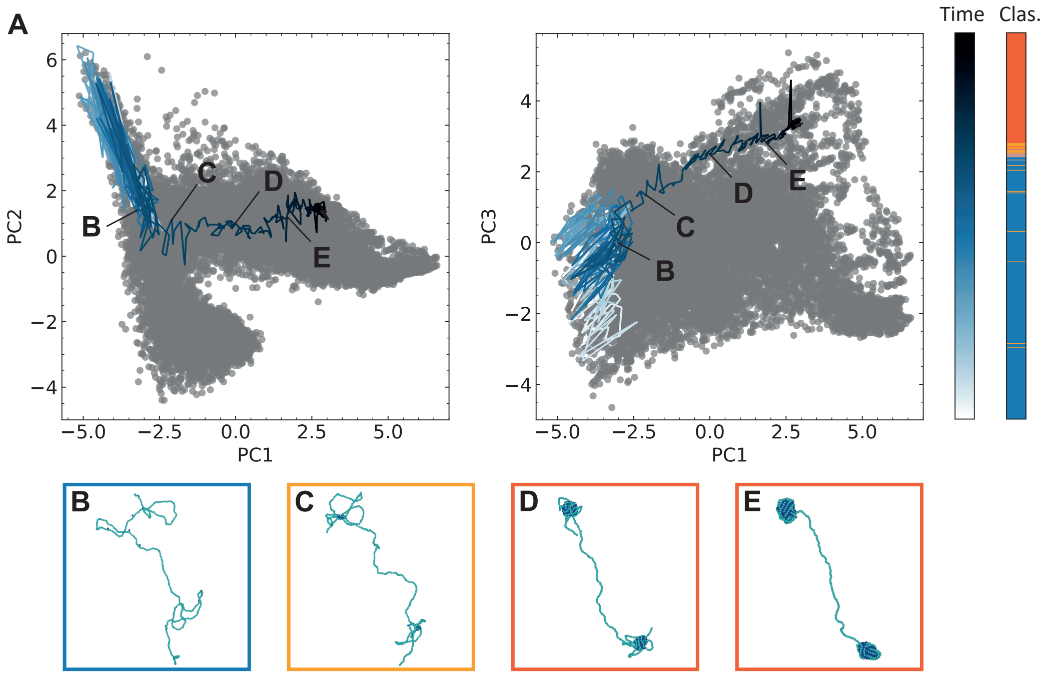


FIG. S6. Typical trajectory in which the protein-induced compaction leads to two condensed DNA-protein complexes. This is for a DNA strand of 408 beads in a system with mutual attractive proteins ($\beta\epsilon_{\mathrm{pp}} = 2.0$) at a concentration of 1450 nM. (*A*) The trajectory on top of the PC1-PC2 and PC1-PC3 distributions. The trajectory is colored with a blue gradient indicating the time (first colorbar) and the second colorbar is the classification according to the GMM (see Fig. 2 of main paper). (*B*-*E*) Show four configurations during the simulation, which locations are indicated in (*A*).
